# Angiocrine FSTL1 insufficiency leads to atrial and vein wall fibrosis via SMAD3 activation

**DOI:** 10.1101/616623

**Authors:** Haijuan Jiang, Luqing Zhang, Xuelian Liu, Wei Sun, Katsuhiro Kato, Chuankai Chen, Xiao Li, Wencan Han, Fujing Zhang, Qi Xiao, Zhongzhou Yang, Zhihai Qin, Ralf H. Adams, Xiang Gao, Yulong He

## Abstract

Angiocrine factors, mediating the endothelial-mural cell interaction in vascular wall construction as well as maintenance, are incompletely characterized. Here we show that loss of follistatin-like protein 1 (FSTL1) in endothelial cells (*Fstl1^ECKO^*) led to an increase of pulmonary vascular resistance, resulting in the heart regurgitation especially with tricuspid valves. However, this abnormality was not detected in mutant mice with *Fstl1* deletion in smooth muscle cells or hematopoietic cells. We further showed that there was excessive alpha-smooth muscle actin (αSMA) associated with atrial endocardia, heart valves, veins and microvessels after the endothelial FSTL1 deletion. Consistently, there was an increase of collagen deposition as demonstrated in livers of *Fstl1^ECKO^* mutants. The SMAD3 phosphorylation was significantly enhanced and pSMAD3 staining was colocalized with αSMA in vein walls, suggesting the activation of TGFβ signaling in vascular mural cells of *Fstl1^ECKO^* mice. The findings imply that endothelial FSTL1 is critical for the homeostasis of atria and veins and its insufficiency may favor cardiovascular fibrosis leading to heart failure.

## Introduction

Vascular endothelial cells (ECs) provide structural support for blood circulation by lining the inner layer of tubular vessels and also actively participate in the assembly of the vascular network during organogenesis by secreting various angiocrine factors (Rafii, Butler et al. 2016). For example, endothelial-derived PDGF-B was required for recruiting mural cells expressing PDGFRβ (Leveen, Pekny et al. 1994, Soriano 1994). Mural cells are essential components of blood vascular walls, including pericytes in microvessels and smooth muscle cells (SMCs) in large blood vessels. They exert important roles in vascular function and homeostasis (Armulik, Genove et al. 2011). TGFβ signaling, a crucial pathway for vascular development, is involved in the regulation of mural cell proliferation and differentiation from mesenchymal cells (Gaengel, Genove et al. 2009). Knockout of the TGFβ pathway-related genes, including TGFβ1, type I or II receptors, endoglin and the downstream SMADs, leads to the defective vascular formation with abnormal recruitment of mural cells (Li, Sorensen et al. 1999, Yang, Castilla et al. 1999, Oh, Seki et al. 2000, Larsson, Goumans et al. 2001, Carvalho, Itoh et al. 2007, Goumans, Liu et al. 2009, Li, Lan et al. 2011, Larrivee, Prahst et al. 2012). On the other hand, TGFβ1 was also shown to be a key mediator in tissue fibrosis, a pathophysiological response of tissues to an insult by the recruitment of inflammatory cells, the activation of myofibroblasts and the deposition of extracellular matrix in the affected tissues (Wynn and Ramalingam 2012, Samarakoon, Overstreet et al. 2013, Harvey, Montezano et al. 2016, Schafer, Viswanathan et al. 2017).

Follistatin-like 1 (FSTL1, also known as TSC36 or FRP) was a secreted glycoprotein first identified as a TGF-β1 inducible factor (Shibanuma, Mashimo et al. 1993). FSTL1 is expressed by many types of cells including endothelial cells and smooth muscle cells (Adams, Larman et al. 2007, Shimano, Ouchi et al. 2011, Miyabe, Ohashi et al. 2014), and exerts biological functions in various tissues in development and diseases (Sylva, Moorman et al. 2013). In the circulatory system, it has been reported that FSTL1 is cardio-protective and epicardial FSTL1 was shown to stimulate the division of pre-existing cardiomyocytes in animal models with myocardial infarction (Lara-Pezzi, Felkin et al. 2008, Oshima, Ouchi et al. 2008, Shimano, Ouchi et al. 2011, Wei, Serpooshan et al. 2015). FSTL1 has also been shown to promote angiogenesis, endothelial cell survival and function (Ouchi, Oshima et al. 2008, Liu, Shen et al. 2010). At the molecular level, FSTL1 was reported to modulate the BMP and TGFβ mediated signaling pathway (Tanaka, Murakami et al. 2010, Geng, Dong et al. 2011, Sylva, Li et al. 2011, Xu, Qi et al. 2012, Prakash, Borreguero et al. 2017).

Although both vascular endothelial cells and smooth muscle cells express high levels of FSTL1, little is known about the differential requirement of specific cell-derived FSTL1 in blood vascular formation and maintenance. We found in this study that deletion of *Fstl1* in endothelial cells, but not in SMCs or blood cells, led to an increase of atria-as well as vein-associated alpha-smooth muscle actin (αSMA) positive cells in several organs examined. Vascular smooth muscle tissues participate in the regulation of peripheral blood pressure via their contraction and relaxation. The pathological vascular remodeling would increase the vascular resistance as reflected by the occurrence of heart regurgitation, leading ultimately to the heart failure of *Fstl1^ECKO^* mice.

## Results

### Tricuspid regurgitation of mutant mice with endothelial *Fstl1* deletion

As mice null for *Fstl1* died at birth (Geng, Dong et al. 2011, Sylva, Li et al. 2011), we generated the mice with *Fstl1* deletion in endothelial cells using the *Tek-Cre* transgenic line (*Fstl1^Flox/−^;Tek-Cre*, named *Fstl1^ECKO^*) (Koni, Joshi et al. 2001). We found that FSTL1 was highly expressed by endothelial cells (Fig. 1A-C). *Fstl1* deletion in endothelial cells was examined by immunostaining for PECAM-1 and FSTL1 (Fig. 1A, arrows), Western blot analysis (Fig. 1B), and quantitative RT-PCR (Fig. 1C). The relative level of *Fstl1* transcripts in mutant lung was reduced to about 35 % of the wildtype control (*Fstl1^Flox/+^*: 1.0 ± 0.27, n=7; *Fstl1^Flox/−^*, 0.57 ± 0.18, n=6; *Fstl1^ECKO^*: 0.35 ± 0.07, n=8). In contrast to the complete *Fstl1* knockout mice, *Fstl1^ECKO^* mice showed lethality starting approximately from the postnatal day 10 (P10). Detailed analysis of the survival rate revealed that about 70% of *Fstl1^ECKO^* mice died by the age of 3-week-old and there was no significant difference between male and female mutants (Supplemental Fig. 1A).

**Fig. 1.**
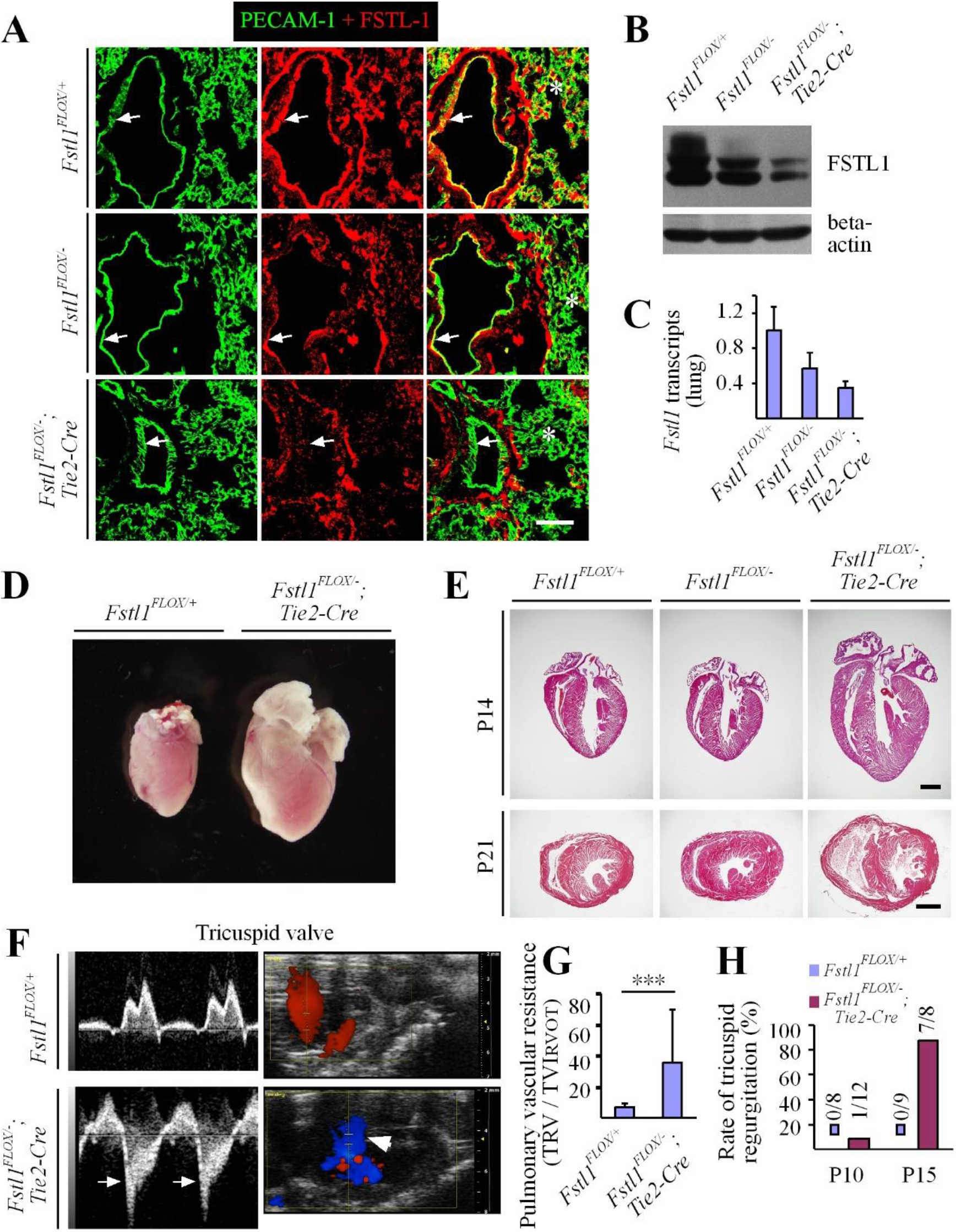
Analysis of FSTL1 expression in endothelial cells and heart regurgitation after the EC-specific deletion of *Fstl1*. **A**. Immunostaining for PECAM-1 and FSTL1 to examine the expression of FSTL1 in blood vascular endothelial cells and its deletion efficiency (green for PECAM-1, and red for FSTL1). Arrows point to endothelium of large vessels and asterisks to the immunostaining signals for FSTL1 in microvessels. **B**. Western blot analysis of FSTL1 in lung of *Fstl1^ECKO^* mutant and control mice. **C**. Quantitative RT-PCR analysis of *Fstl1* transcripts in lung from *Fstl1^Flox/−^;Tek-cre* (*Fstl1^ECKO^*) and littermate control mice. **D, E**. Cardiac hypertrophy in *Fstl1^ECKO^* mice at the postnatal day 21 (P21, D), and histological analysis of hearts from *Fstl1^ECKO^* and control mice at the postnatal day 14 (P14) and P21 (E). **F, G**. Analysis of heart function by *In vivo* ultrasonic imaging (F, P15-17) and quantification of pulmonary vascular resistance in *Fstl1^ECKO^* mice (G, P15). Blue signals in F indicate the backflow of blood in the right ventricle (arrows). **H**. Rate of tricuspid regurgitation at P10 and P15. Note that the tricuspid regurgitation was rare at P10 but occurred in most *Fstl1* mutant mice at P15 (G). Scale bar: 50 μm in A; 1000 μm in E.

Further analysis revealed that the hearts of *Fstl1^ECKO^* mice became enlarged (Fig. 1D), as also demonstrated by the histological analysis at both P14 and P21 (Fig. 1E) and the functional analysis at P10 and P15-17 by the ultrasound imaging system (VisualSonics Vevo 2100, Fig. 1F-H). There was a significant increase of heart to body weight (HBW) ratio in *Fstl1^ECKO^* mice compared with littermate controls (*Fstl1^Flox/+^*: 0.79 ± 0.13 %, n=10; *Fstl1^Flox/−^*: 0.78 ± 0.06 %, n=8; *Fstl1^ECKO^*: 1.48 ± 0.55 %, n=7; *Fstl1^ECKO^* vs *Fstl1^Flox/+^*:: P=0.0014). The pulmonary vascular resistance (PVR) was calculated by TRV (tricuspid regurgitant velocity) / TVI_RVOT_ (right ventricular outfow time-velocity integral). There was a significant increase of PVR in *Fstl1^ECKO^* mice at P15 (Fig. 1G; *Fstl1^Flox/+^*: 6.58 ± 2.85, n=9; *Fstl1^ECKO^*: 35.56 ± 34.15, n=7; P=0.0227). The alteration of vascular resistance was also demonstrated by the observation that most of the mutant mice (7 out of 8 mice) showed tricuspid regurgitation at P15 (Fig. 1F), while there was only 1 out of 12 *Fstl1^ECKO^* mice detected at P10 (Fig. 1H). However, only 1 out of 5 *Fstl1^ECKO^* mice showed the mitral regurgitation while 5 out of 5 displayed the tricuspid regurgitation. Consistently, there was a significant increase of the right ventricle internal diameter (diastole, Supplemental Figure 1B; *Fstl1^Flox/+^*: 0.91 ± 0.35, n=9; *Fstl1^ECKO^*: 1.35 ± 0.39, n=9; P=0.0244) and also the right ventricular cavity area (diastole, Supplemental Figure 1C; *Fstl1^Flox/+^*: 3.43 ± 0.73,n=9; *Fstl1^ECKO^*: 5.92 ± 2.14, n=9; P=0.0044). There may be a trend of increase but no significant difference was detected with the left ventricule internal diameter (*Fstl1^Flox/+^*: 2.92 ± 0.17, n=9; *Fstl1^ECKO^*: 3.20 ± 0.48, n=9; P=0.1159) and the left ventricular cavity area (*Fstl1^Flox/+^*: 6.15 ± 0.47, n=9; *Fstl1^ECKO^*: 7.69 ± 2.52, n=9; P=0.0894).

### Increase of αSMA associated with atrial endocardia and valves of *Fstl1^ECKO^* mice

To investigate causes of the lethality with *Fstl1^ECKO^* mice, we examined heart by immunostaining for αSMA and PECAM-1. There was a dramatic increase of endocardium-associated αSMA^+^ cells in right and left atria (Fig. 2A, B). Microvessels in the heart walls of *Fstl1* mutant mice also showed an increase of αSMA staining, which is rarely detected in those of control mice (Fig. 2A, B). Consistently, a similar change of the αSMA staining, associated with the endothelial cells of valve leaflets, was also detected in the tricuspid and mitral valves (Fig. 2C, D). However, there was no αSMA staining associated with endocardium of right and left ventricles, and there was also no obvious change of αSMA^+^ microvessels detected in the ventricular walls (Supplemental Fig. 2).

**Fig. 2.**
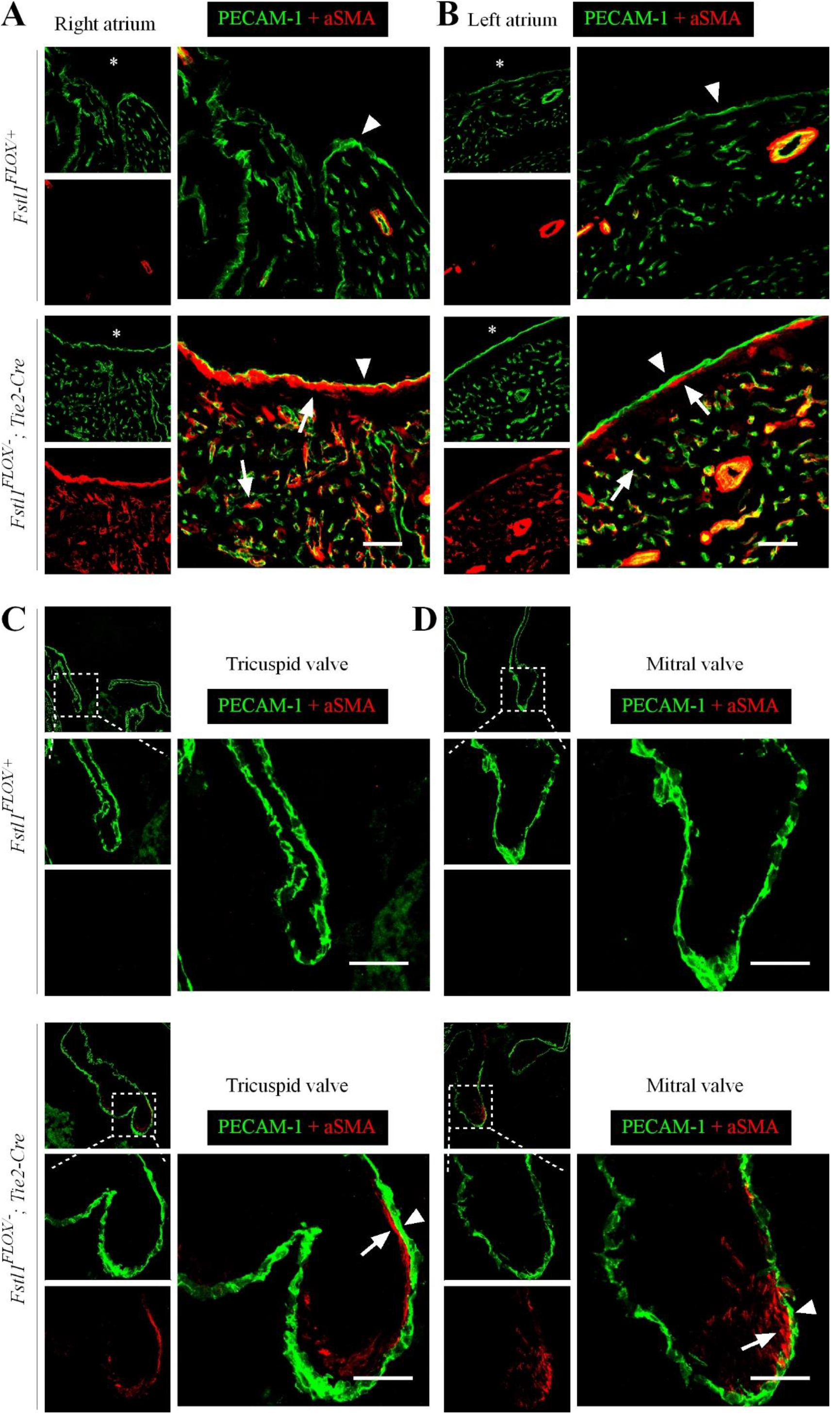
Increased aSMA association with the atrial endocardium and heart valves of *Fstl1^ECKO^* mutants. **A, B**. Immunostaining analysis of blood vessels for PECAM-1 (green) and αSMA (red) with right atrium (A), left atrium (B) of *Fstl1^ECKO^* mice (P18). Asterisks indicate heart cavity, and arrows point to abnormal association of αSMA with endocardium (arrowheads). **C, D**. Analysis of tricuspid and mitral valves from *Fstl1^ECKO^* and littermate control mice by immunostaining (P18). Arrows point to the αSMA association of with valve endothelia (arrowheads). Scale bar: A-D, 50 μm.

To further investigate the effect of non-EC-derived FSTL1 in vascular development, we generated the doubly transgenic mice with *Fstl1* deletion in smooth muscle cells (*Fstl1^Flox/−^;αSMA-Cre*, named *Fstl1^SMCKO^*) or in hematopoietic cells (*Fstl1^Flox/−^;Vav-iCre*, named *Fstl1^HCKO^*). FSTL1 deletion in *Fstl1^SMCKO^* mice was examined by Western blot analysis (Supplemental Fig. 3A), and quantitative RT-PCR (*Fstl1^Flox/+^*: 1.0 ± 0.17, n=5; *Fstl1^Flox/−^*: 0.46 ± 0.18, n=6; *Fstl1^SMCKO^*: 0.30 ± 0.07, n=7). Deletion of *Fstl1* gene in hematopoietic cells of *Fstl1^HCKO^* mice was confirmed by the PCR genotyping of blood cells from mutant or control mice (Supplemental Fig. 3B). All the *Fstl1^SMCKO^* and *Fstl1^HCKO^* mutant mice survived well by the age of 3-week-old. Surprisingly, although SMCs are one of the major sources of FSTL1 and the SMA-Cre mediated deletion efficiency of FSTL1 is similar to that of *Fstl1^ECKO^* mice at the stage. There were no obvious defects observed in mutant hearts (Supplemental Fig. 3C) or in the smooth muscle cell coverage with blood vessels of *Fstl1^SMCKO^* mice (Supplemental Fig. 3E). Also, there were no detectable heart and vascular abnormalities in *Fstl1^Flox/−^;Vav-iCre* mice (Supplemental Fig. 3D and F).

### Excessive αSMA association with lung veins and microvessels in FSTL1 mutant mice

Consistent with the observation in the atria, there was an obvious increase of αSMA staining with blood vessels of lung in *Fstl1^ECKO^* mice (Fig. 3A). The alteration of αSMA expression was further validated by Western blot analysis in lung (Fig. 3B). The ratio of αSMA to β-actin was quantified and normalized against the control group (Fig. 3C). There was a significant increase of αSMA in *Fstl1^ECKO^* mice compared with that of the littermate controls (Fig. 3C; *Fstl1^Flox/+^*: 1.0 ± 0.11, n=7; *Fstl1^Flox/−^*: 0.95 ± 0.22, n=6; *Fstl1^ECKO^*: 1.32 ± 0.24, n=9; *Fstl1^ECKO^* vs *Fstl1^Flox/+^*: P=0.0060). Interestingly, endomucin, a membrane-bound glycoprotein expressed luminally by venous and capillary ECs, but not by arterial endothelium, was significantly decreased in lungs of the mutant mice (Fig. 3C; *Fstl1^Flox/+^*: 1.0 ± 0.05, n=7; *Fstl1^Flox/−^*: 0.95 ± 0.12, n=6; *Fstl1^ECKO^*: 0.70 ± 0.19, n=9; *Fstl1^ECKO^* vs *Fstl1^Flox/+^*: P=0.0010). The decrease of endomucin level could also reflect the alteration of blood pressure (Zahr, Alcaide et al. 2016). Interestingly, we found that the increased αSMA was mainly associated with veins and vein-associated microvessels, which were positive for the venous and capillary endothelial cell marker endomucin (Fig. 3A, arrows; lung). This was further validated by the immunostaining for Dll4, an arterial endothelial cell marker. The blood vessels with abnormal αSMA coating were negative for Dll4 (Fig. 3D, arrowheads; lung).

**Fig. 3.**
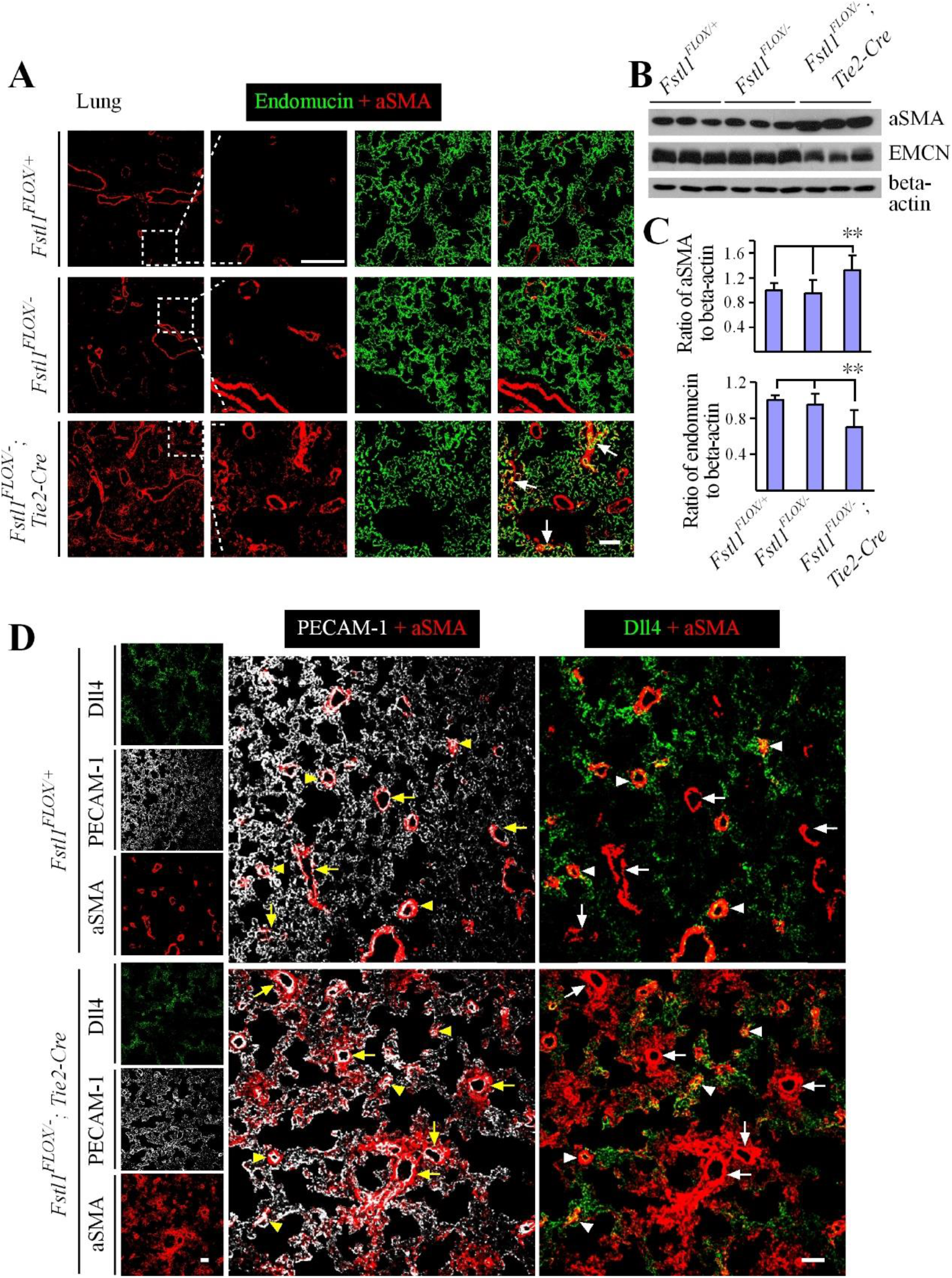
Upregulation of αSMA in blood vessels negative for the arterial marker Dll4 in lung of *Fstl1^ECKO^* mice. **A**. Immunostaining analysis of blood vessels for endomucin (green) and αSMA (red) with lung from *Fstl1^ECKO^* and control mice (P21). Arrows point to the αSMA staining in big and microvessels. **B, C**. Western blot analysis of αSMA and endomucin expression in lung from *Fstl1^ECKO^* and littermate control mice (B), and quantification of the ratio of αSMA or endomucin to beta-actin respectively (C). **D**. Immunostaining analysis of blood vessels for Dll4 (green), αSMA (red), and PECAM-1 (white) with lung from *Fstl1^ECKO^* and control mice. Arrows point to veins (Dll4 negative) and arteries (arrowheads, Dll4^+^). Note that there is a massive increase of αSMA association with veins and microvessels, but this is not obvious with arteries at the same stage. Scale bar: 50 μm in A and D.

### Occurrence of blood vessel-associated fibrosis in livers of FSTL1 mutant mice

We further analyzed the vascular associated αSMA in several other tissues including liver, kidney, retina and skin. The increase of αSMA staining was detected by immunostaining in liver (Fig. 4A), and this was confirmed by Western blot analysis (Fig. 4B and C; *Fstl1^Flox/+^*: 1.0 ± 0.48, n=6; *Fstl1^Flox/−^*: 0.84 ± 0.10, n=3; *Fstl1^ECKO^*: 3.45 ± 2.17, n=6; *Fstl1^ECKO^* vs *Fstl1^Flox/+^*: P=0.0220). Consistent with the results observed in lung, the protein level of endomucin was significantly decreased (Fig. 4D; *Fstl1^Flox/+^*: 1.0 ± 0.30, n=6; *Fstl1^Flox/−^*: 0.83 ± 0.36, n=6; *Fstl1^ECKO^*: 0.39 ± 0.14, n=6; *Fstl1^ECKO^* vs *Fstl1^Flox/+^*: P=0.0011). An obvious increase of αSMA was also detected in kidney (Supplemental Fig. 4). It is worth pointing out that there is variation about the vascular phenotype among organs of *Fstl1^ECKO^* mice by the age of 3-week-old, with the increase of αSMA^+^ in 14 out of 14 (14/14) livers, 10/14 lungs and 5/11 kidneys examined. However, there was no obvious difference in αSMA associated with blood vessels in skin (Supplemental Fig. 5A) and retina tissues examined by P21 (Supplemental Fig. 5B). FSTL1 deficiency in endothelial cell did not produce any obvious effect on lymphatic vessel development as shown in trachea (Supplemental Fig. 5C). This suggests that there may be a differential requirement of endothelial FSTL1 in the regulation of organ-specific vascular systems.

**Fig. 4.**
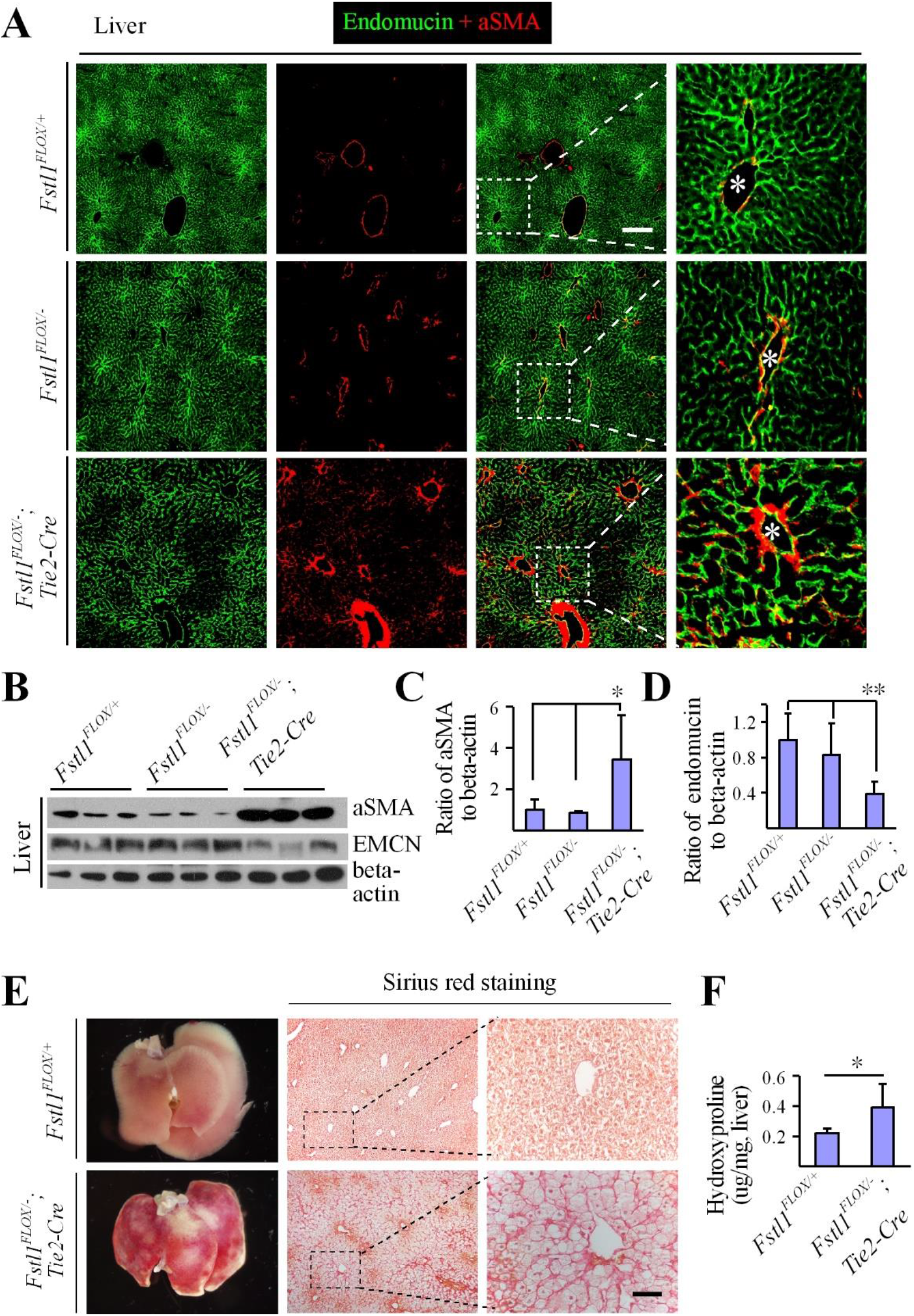
Occurrence of vascular fibrosis in liver of *Fstl1^ECKO^* mutant mice. **A**. Immunostaining analysis of blood vessels for endomucin (green) and αSMA (red) with liver from *Fstl1^ECKO^* mutant and littermate control mice (P21). Asterisks point to the veins. **B**. Analysis of αSMA and endomucin expression in liver from *Fstl1^ECKO^* and control mice by Western blot analysis. **C, D**. Quantification of the ratio of αSMA or endomucin to β-actin respectively. **E, F**. Analysis of liver fibrosis by Sirius red staining (E) and quantification of hydroxyproline (F). Scale bars: 200 μm in A, 50 μm in E.

Consistent with the increase of blood vessel-associated αSMA, we also detected the increase of collagen deposition by Sirius red staining in liver of *Fstl1^ECKO^* mice (Fig. 4E). This was confirmed by the quantification of hydroxyproline content in liver tissue lysates (Fig. 4F). There was a significant increase of hydroxyproline in *Fstl1* mutant mice compared with the littermate control (*Fstl1^Flox/+^*: 0.22 ± 0.03 μg / mg, n=6; *Fstl1^ECKO^*: 0.39 ± 0.16 μg / mg, n=9; P=0.0253).

### SMAD3 activation in vein walls of *Fstl1^ECKO^* mutant mice

To check whether there is any transition of endothelial cells into αSMA^+^ cells in blood vessels, we introduced ROSA^mT/mG^ allele into the *Fstl1^Flox/−^;Tek-cre* line and the *Fstl1^Flox/+^;Tek-cre;mT/mG* mice was used as control. Interestingly, we found that the αSMA staining was detected mainly at the outer layer of vessel walls in both large and microvessels in the liver of *Fstl1^Flox/−^;Tek-cre;mT/mG* mutant mice (Fig. 5A). This suggests that the upregulation of αSMA occurred in peri-vascular mural cells. Consistent with the observations in lung, we found that the increased αSMA was associated with veins and microvessels, which were positive for the venous endothelial cell marker EphB4 (Fig. 5B, arrows; arrowheads point to arteries).

**Fig. 5.**
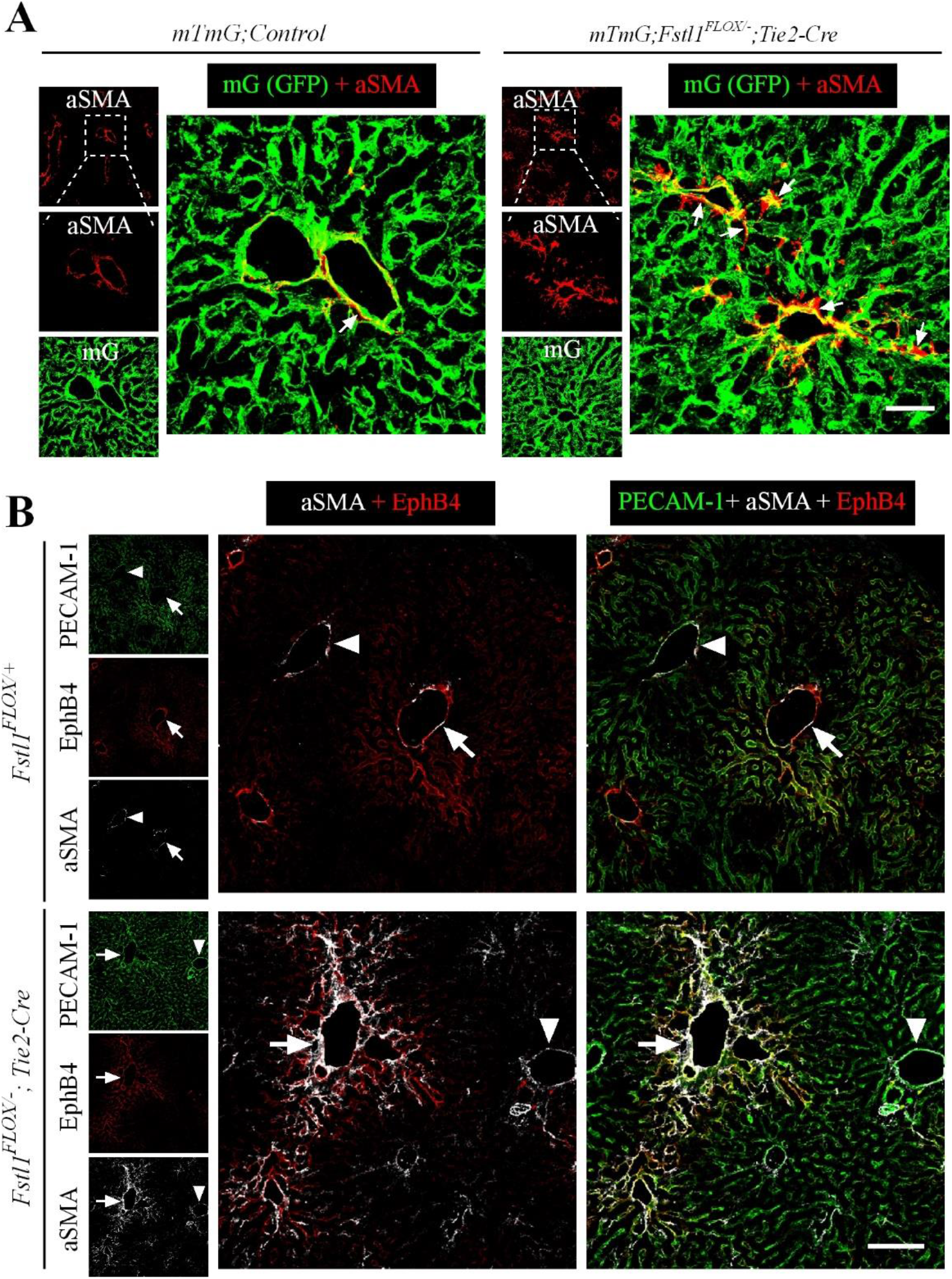
Association of αSMA with hepatic veins and microvessels positive for the venous marker EphB4 after the endothelial FSTL1 deletion. **A**. Analysis of smooth muscle cell coverage (αSMA^+^) with blood vessels (GFP^+^) in *mTmG; Fstl1^ECKO^* and littermate control mice at P18. Arrows point to the large and microvessels positive for αSMA staining. **B**. Analysis of blood vessels in liver from *Fstl1^ECKO^* and control mice by immunostaining for PECAM-1 (green), EphB4 (red) and αSMA (white). Note that αSMA mainly associated with EphB4 positive vessels (veins, arrows; arrowheads to arteries) in liver of *Fstl1^ECKO^* mice. Scale bars: 100 μm in A and B.

To examine the alteration of TGFβ mediated signaling, we analyzed the phosphorylation of SMAD3 by the immunostaining (Fig. 6A, B) and western blot analysis (Fig. 6C, D). The number of pSMAD3^+^ /αSMA^+^ doubly positive cells associated with blood vessels was significantly increased in livers of *Fstl1^ECKO^* mice compared with that of littermate controls (Fig. 6A and B; *Fstl1^Flox/+^*: 1.3 ± 0.2 / grid, n=4; *Fstl1^ECKO^*: 7.3 ± 1.1 / grid, n=4; P=0.0001). This was further confirmed by Western blot analysis and the quantification of pSMAD3: tSMAD2/3 in liver (the ratio normalized against the control as shown in Fig. 6C and D; *Fstl1^Flox/+^*: 1.0 ± 0.14, n=6; *Fstl1^ECKO^*: 1.83 ± 0.46, n=7; P=0.0014). There was also a slight increase detected with the ratio of pSMAD1/5/8: tSMAD1 (Fig. 6D; *Fstl1^Flox/+^*: 1.0 ± 0.06, n=6; *Fstl1^ECKO^*: 1.14 ± 0.14, n=6; P=0.0425). A similar increase of SMAD3 activation was also shown in lung (pSMAD3 /tSMAD2/3: *Fstl1^Flox/+^*: 1.0 ± 0.23, n=6; *Fstl1^ECKO^*: 1.44 ± 0.26, n=7; P=0.0086), while there was no significant difference observed with pSMAD1/5/8: tSMAD1 (*Fstl1^Flox/+^*: 1.0 ± 0.31, n=6; *Fstl1^ECKO^*: 1.08 ± 0.40, n=6; P=0.7047).

**Fig. 6.**
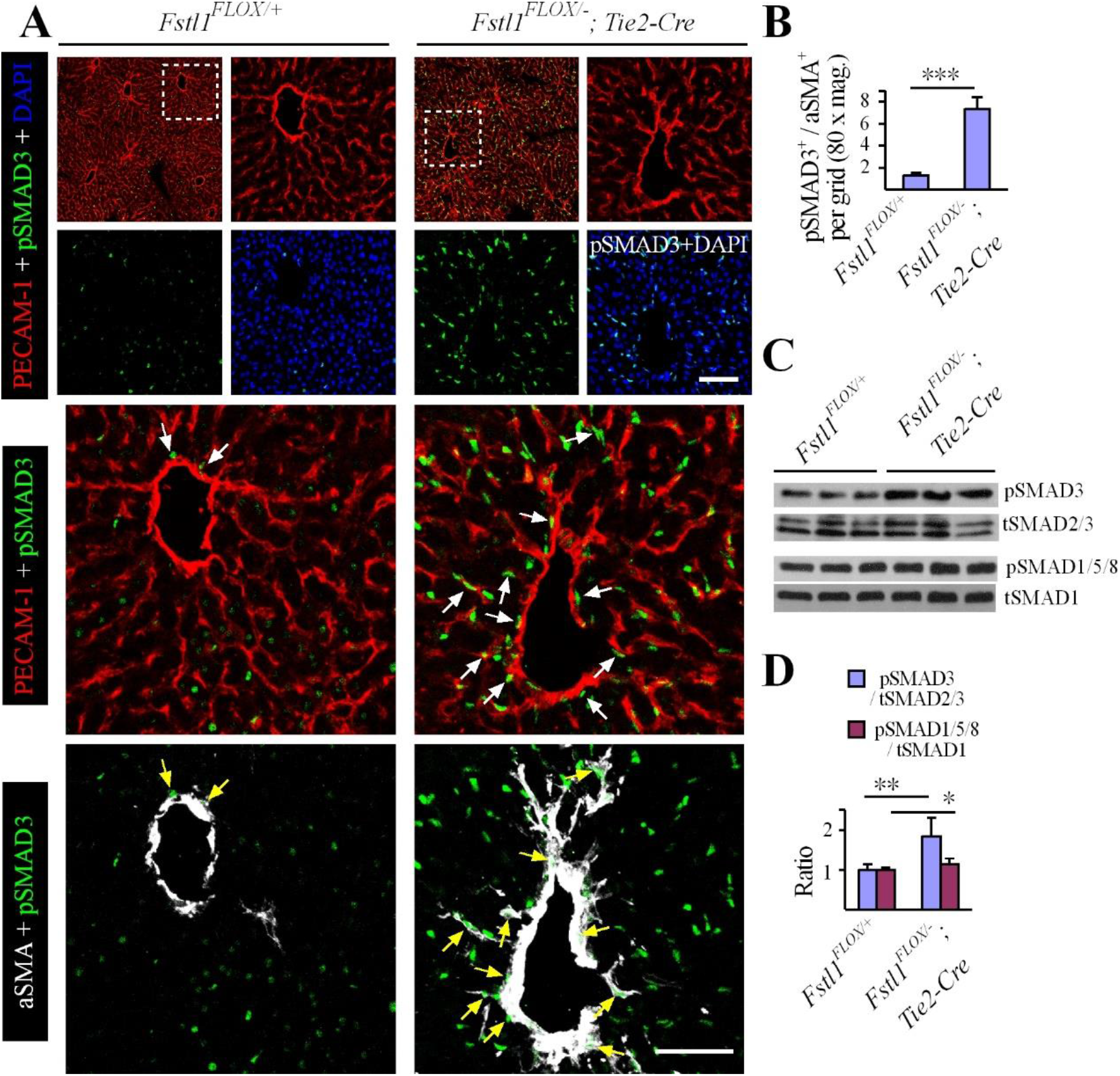
SMAD3 activation in vascular mural cells of *Fstl1^ECKO^* mutant mice. **A, B**. Analysis of pSMAD3 positive cells by immunostaining and the quantification of pSMAD3^+^ /αSMA^+^ doubly positive cells (arrows) surrounding blood vessels in *Fstl1^ECKO^* mutants compared with the littermate control mice (P18). **C, D**. Western blot analysis of pSMAD3 and pSMAD1/5/8 in liver tissues (C) and the quantification of the ratio of phosphorylated and total proteins was shown in D. Scale bar: 50 μm in A.

## Discussion

We have shown in this study that the EC-derived FSTL1 is required for the homeostasis of cardiovascular system. Lack of endothelial FSTL1 led to a dramatic increase of αSMA association with veins and associated microvessels, in addition to the abnormal remodeling with atrial endocardia as well as heart valves. At the molecular level, an enhanced activation of SMAD3 signaling was detected in vascular mural cells in the absence of endothelial FSTL1, while the SMAD1/5/8 pathway was only mildly affected. The excessive αSMA would promote the cardiac rigidity and decrease the vascular elasticity. This may directly account for the alteration of vascular resistance as reflected by the occurrence of tricuspid regurgitation and the heart failure of the EC-specific *Fstl1* mutant mice.

It was previously reported that deletion of FSTL1 led to the abnormal development of mitral valves (Prakash, Borreguero et al. 2017). It is worth noting that the authors used a different Tie2-Cre deletor from the line used in our study (Koni, Joshi et al. 2001). With the Tie2-Cre transgenic mice used by Prakash et al. (Prakash, Borreguero et al. 2017), the Cre recombinase-mediated gene recombination occurred not only in endothelial cells, but also in mesenchymal cells of atrioventricular canal and the proximal cardiac outflow tract (Kisanuki, Hammer et al. 2001). It is likely that the defects with mitral valves observed by Prakash et al. was at least partly due to the loss of FSTL1 in mesenchymal cells (Prakash, Borreguero et al. 2017). In this study, we found that the tricuspid valves displayed a more severe abnormality with the regurgitation detected earlier than that of mitral valves. We speculate that the lethality of *Fstl1^ECKO^* mice in our study may result from the abnormal remodeling of cardiovascular system in a systemic level. Firstly, the excessive αSMA^+^ cell investment with vessel walls detected in several organs would increase the peripheral vascular resistance. Secondly, the increased association of αSMA with atrial endocardia and heart valves would increase the cardiac rigidity. Both factors would ultimately contribute to the hypertension and heart failure of *Fstl1^ECKO^* mutant mice.

Pathogenesis of pulmonary hypertension often involves the abnormal vascular muscularization affecting mainly arterioles, such as the increased proliferation of vascular SMCs (Voelkel, Natarajan et al. 2011, Tuder, Archer et al. 2013). It has been shown that TGFβ mediated signaling is implicated in the development of pulmonary hypertension, with genetic mutations identified in several genes of this pathway including ALK1, BMPR2, endoglin, BMP9 and the downstream mediators (Goumans, Liu et al. 2009, Machado, Southgate et al. 2015, Wang, Fan et al. 2016). BMPR2 haploinsufficiency was shown to cause pulmonary arterial hypertension (PAH) (Machado, Pauciulo et al. 2001), while overexpression of BMPR2 in endothelial cells ameliorated PAH via the suppression of Smad2/3 signaling (Harper, Reynolds et al. 2016). Furthermore, targeted disruption of SMAD3 signaling suppressed the pathological processes of vascular or interstitial fibrosis in several other organs examined (Sato, Muragaki et al. 2003, Wang, Huang et al. 2006, Lee, Wright et al. 2016, Liu, Das et al. 2017). FSTL1 is a TGFβ inducible factor (Shibanuma, Mashimo et al. 1993). We found in this study that the endothelial FSTL1 deficiency led to a significant increase of SMAD3 activation but only a mild change with SMAD1/5/8 phosphorylation, suggesting a negative effect of FSTL1 on TGFβ pathway. Consistently, we observed that there was an upregulation of αSMA expression in the cardiovascular system, accounting for the occurrence of hypertension in *Fstl1^ECKO^* mutants. The alteration of vascular resistance in *Fstl1^ECKO^* mice was also reflected by the decrease of luminally expressed endomucin in venous and capillary endothelia, as fluid shear stress was shown to downregulate its surface localization (Zahr, Alcaide et al. 2016). Surprisingly, we found in this study that the increase of αSMA mainly occurred with atria and vein-associated vessels, which was accompanied by the increased collagen deposition as demonstrated in liver, a typical feature of tissue fibrosis. It is possible that EC-derived FSTL1 may maintain the quiescence of peri-endothelium / -endocardium mural cells or mesenchymal cells by antagonizing TGFβ signaling. It remains to be investigated about the mechanism underlying the discrepancy between veins versus arteries in response to the loss of endothelial FSTL1. In addition, it is unclear why the abnormal vascular remodeling occurred postnatally with *Fstl1^ECKO^* mice, but without any obvious abnormalities observed during embryogenesis even in *Fstl1* conventional knockout mice (our unpublished observation).

In addition, FSTL1 is expressed also by vascular mural cells and the muscle-derived FSTL1 was shown to have beneficial effects on cardiovascular functions or tissue repair in pathological conditions such as pressure overload or injury (Shimano, Ouchi et al. 2011, Miyabe, Ohashi et al. 2014). However, we found that deletion of FSTL1 in SMCs did not affect blood vascular formation and remodeling in development. This is consistent with the recent finding that FSTL1 from different cell-types (e.g. myocardial versus epicardial cells) exerts distinct functions in heart, which may be due to the differential protein glycosylation (Wei, Serpooshan et al. 2015). Furthermore, the significant increase of αSMA was detected in atria, liver, lung and kidney of *Fstl1^ECKO^* mice, but not obvious with skin and retinal blood vessels examined. This implies a potential role of tissue microenvironment on the biological functions of FSTL1.

To summarize, the requirement of EC-derived FSTL1 for vascular homeostasis adds a new factor to the regulatory network for vascular health. It is still to be investigated about the differential effects of EC- versus SMC-derived FSTL1 on vascular maintenance and the distinct requirement of endothelial FSTL1 in different organs. Furthermore, this study also introduced a genetically modified disease model for the investigation of mechanisms underlying pathological heart remodeling related to pulmonary or systemic hypertension.

## Methods

### Animal work

All animal experiments were performed in accordance with the institutional guidelines of Soochow University Animal Center. Conditional knockout mice targeting *Fstl1* gene (*Fstl1^Flox^*) were generated as previously described (Li, Zhang et al. 2011). *Fstl1^+/−^* mice were obtained by crossing of *Fstl1^Flox^* mice with EIIA-Cre mice (Lakso, Pichel et al. 1996). *Tek-Cre* mice were used to generate the endothelial cell specific knockout mouse models (*Fstl1^Flox/−;Tek-Cre^*, named *Fstl1^ECKO^*) (Koni, Joshi et al. 2001). For the lineage tracing, we introduced *ROSA^mT/mG^* (The Jackson Laboratory, Stock No. 007576) allele into the *Fstl1* mutant mice. αSMA-Cre and Vav-iCre mouse lines were used for *Fstl1* deletion in smooth muscle cell (*Fstl1^Flox/−^;αSMA-Cre*, named *Fstl1^SMCKO^*) or hematopoietic cell (*Fstl1^Flox/−^;Vav-iCre*, named *Fstl1^HCKO^*) (de Boer, Williams et al. 2003, Wu, Yang et al. 2007). To examine the floxed allele, Forward primer CTCCCACCTTCGCCTCTAAC and Reverse primer ATTTCTGCTCCTAGCGTGCC were used to amplify a 400 bp fragment for the wild type (WT) allele and a 506 bp fragment for the floxed allele. For the genotyping of *Fstl1^+/−^* mice, primers used are as follows: Forward primer CTCCCACCTTCGCCTCTAAC, Reverse primer CGGCTAGGAAAGACTTGGAA. The WT allele gives a product of 635 bp and the knockout allele gives a band of 341 bp. Cre mice were genotyped using the following primers (Forward CAACGAGTGATGAGGTTCGCAAG and Reverse TGATCCTGGCAATTTCGGCTATAC).

### Echocardiography

Echocardiography of FSTL1 mutant and control mice (P10-P17) was performed using a Vevo 2100 ultrasound system (VisualSonics). Mice were anesthetized with isoflurane at 3 % for induction and at 1-1.5 % for maintenance. Skin hairs of chest regions were removed. For the analysis of heart function, peak tricuspid regurgitation velocity (TRV) was measured using pulsed-wave Doppler and the highest velocity recorded from multiple views was used, including apical 4-chamber, parasternal and subcostal views.

### Immunostaining

For whole mount immunostaining, ears were harvested, fixed in 4 % paraformaldehyde, blocked with 3 % (w/v) milk in PBS-TX (0.3 % Triton X-100), and incubated with primary antibodies overnight at 4 °C. For the frozen tissue section staining, tissues were collected and fixed in 4 % PFA for 2 h at 4°C, followed by the incubation in 20% sucrose overnight before being embedded in OCT. Frozen sections of 10 μm were used. The antibodies used were rat anti-mouse PECAM-1 (BD Pharmigen, 553370), Cy3-αSMA (Sigma C6198), Endomucin (eBioscience, 14-5851), pSMAD3 (abcam ab52903) and FSTL1 (R&D, AF1738). Appropriate Alexa 488, Alexa594 (Invitrogen) conjugated secondary antibodies were used. All fluorescently labeled samples were mounted and analyzed with a confocal microscope (Olympus Flueview 1000), or Leica MZ16F fluorescent dissection microscope.

### Quantitative real-time RT-PCR

Lung tissues from *Fstl1* mutant and control mice were collected and homogenized in Trizol (Ambion). RNA extraction and reverse transcription was performed following standard procedures (RevertAid First Strand cDNA Synthesis Kit, Thermo Scientific). Quantitative real-time RT–PCR was carried out using the SYBR premix Ex Taq kit (TaKaRa). Briefly, for each reaction, 50 ng of total RNA was transcribed for 2 min at 50°C and a denaturing step at 95°C for 30 s, then followed by 40 cycles of 5 s at 95°C and 34 s at 60°C. Fluorescence data were collected and analyzed using ABI PRISM 7500. The primers used were as follows: FSTL1: 5’-TCTGTGCCAATGTGTTTTGTGG-3’, 5’-TGAGGTAGGTCTTGCCATTACTG-3’; GAPDH: 5’-GGTGAAGGTCGGTGTGAACG-3’, 5’-CTCGCTCCTGGAAGATGGTG-3’.

### Sirius red staining and hydroxyproline quantification

Collagen deposition in tissues were analyzed by sirius red staining and the quantification of hydroxyproline. Briefly, tissues were collected and fixed in 4% paraformaldehyde overnight at 4°C. Paraffin sections (6 μm) were processed for staining first with hematoxylin, followed by the Picro-sirius red (Sigma-Aldrich) staining for one hour (0.5 g Direct Red 80 dissolved in 500 ml 1.3% Picric acid from Sangon, Shanghai). Sections were then washed, dehydrated and mounted in the mounting medium for further microscopic analysis. For the measurement of tissue collagen content by the hydroxyproline quantification, liver tissues (60 mg) were subjected to the alkaline hydrolysis and hydroxyproline concentration in the tissue lysate was determined following the manufacturer’s instructions (Jiancheng, Nanjing, China) as follows: (Sample OD value – blank OD) / (standard OD–blank OD) × standard sample concentration (5 μg / ml) × total hydrolysate volume (10 ml) / tissue wet weight (mg).

### Western blot

For the analysis of FSTL1 expression and deletion efficiency, tissues were washed with ice cold PBS and lysed in lysis buffer (1 mM PMSF, 2 mM Na_3_VO_4_, 1× protease and phosphatase inhibitor cocktail tablets without EDTA from Roche Applied Science, 20 mM Tris-HCl pH 8.0, 100 mM NaCl, 10 % Glycerol, 50 mM NaF, 10 mM β-glycerolphosphate, 5 mM Sodium pyrophosphate, 5 mM EDTA, 0.5 mM EGTA, 1 % NP-40). The lysates were incubated on ice for 0.5 hour with rotation and was clarified by centrifugation. Protein concentration was determined using the BCA protein assay kit (PIERCE). Equal amounts of protein were used for analysis. Antibodies used include goat anti-mouse FSTL1 (R&D, BAF1738), mouse anti-all αSMA (sigma A2547), pSMAD3 (abcam ab52903), tSMAD2/3 (R&D3797), pSMAD1/5/8 (cell signaling 9511S), tSMAD1 (cell signaling 9743S), and mouse monoclonal to β–actin antibody (C4, Santa Cruz sc-47778).

### Statistics

Statistical analysis was performed with the unpaired t test. All statistical tests were two-sided.

## Acknowledgement

We thank Drs. Rodrigo Dieguez-Hurtado, Cong Xu and Qi Chen of Max-Planck-Institute for Molecular Biomedicine for helpful discussion about the project, and staff in the Animal facility of Soochow University and Model Animal Research Institute of Nanjing University for technical assistance. This work was supported by grants from the National Natural Science Foundation of China (91539101, 81770489, 91739304), the Key Program of Natural Science Foundation of Jiangsu Higher Education Institutions (18KJA180012) and the Priority Academic Program Development of Jiangsu Higher Education Institutions.

## Author contributions

HJ, LZ, XL: acquisition and analysis of data, revising the manuscript and final approval of the version; WS, KK, CC, XL, WH, FZ, QX: analysis of data, revising the manuscript and final approval of the version; ZY, ZQ, RHA, XG: interpretation of data, revising the manuscript and final approval of the version; YH: conception and design of the work, analysis and interpretation of data, drafting the manuscript and final approval of the version.

## Conflict of interest

None

